# *Brassica rapa* domestication: untangling wild and feral forms and convergence of crop morphotypes

**DOI:** 10.1101/2021.04.05.438488

**Authors:** Alex C. McAlvay, Aaron P. Ragsdale, Makenzie E. Mabry, Xinshuai Qi, Kevin A. Bird, Pablo Velasco, Hong An, J. Chris Pires, Eve Emshwiller

## Abstract

The study of domestication contributes to our knowledge of evolution and crop genetic resources. Human selection has shaped wild *Brassica rapa* into diverse turnip, leafy, and oilseed crops. Despite its worldwide economic importance and potential as a model for understanding diversification under domestication, insights into the number of domestication events and initial crop(s) domesticated in *B. rapa* have been limited due to a lack of clarity about the wild or feral status of conspecific non-crop relatives. To address this gap and reconstruct the domestication history of *B. rapa*, we analyzed 68,468 genotyping-by-sequencing-derived SNPs for 416 samples in the largest diversity panel of domesticated and weedy *B. rapa* to date. To further understand the center of origin, we modeled the potential range of wild *B. rapa* during the mid-Holocene. Our analyses of genetic diversity across *B. rapa* morphotypes suggest that non-crop samples from the Caucasus, Siberia, and Italy may be truly wild, while those occurring in the Americas and much of Europe are feral. Clustering, tree-based analyses, and parameterized demographic inference further indicate that turnips were likely the first crop type domesticated, from which leafy types in East Asia and Europe were selected from distinct lineages. These findings clarify the domestication history and nature of wild crop genetic resources for *B. rapa*, which provides the first step toward investigating cases of possible parallel selection, the domestication and feralization syndrome, and novel germplasm for *Brassica* crop improvement.

## Introduction

Domestication is a process of adaptation to agroecological niches and human preferences (Larson et al., 2014) driven by a complex mix of ecological, biological, and cultural factors (Price et al., 2011; Gepts et al., 2012). The study of domestication provides insight into the nature of contemporary crop genetic resources (Gepts et al., 2012; Zeder, 2015) and evolutionary processes in general (Darwin, 1868; Fuller et al., 2014). The identity of the wild relatives of a crop is a key piece of information in disentangling the often-complex domestication history, allows the direct comparison of wild to domesticated forms (Page et al., 2019), and can contribute useful alleles responsible for agronomically important traits (Zhang et al., 2017). However, the identification of truly wild populations is often complicated by the presence of conspecific feral populations which are common for many crops (e.g., Wang et al. 2017). Unlike wild populations, feral plants derive from domesticated crops that have either escaped from cultivation on their own (endoferal) or through crossing with wild populations (exoferal; Gressel, 2005). Additionally, complex histories of hybridization and introgression among wild, feral and domesticated forms within and across species make identification of truly wild populations difficult (Beebe et al., 1997; Wang et al., 2017; Allaby et al., 2008).

The morphologically diverse crops in the genus *Brassica* (e.g., broccoli, cabbage, bok choy, and canola) provide powerful study systems to understand domestication and artificial selection (Gómez-Campo and Prakash, 1999; Bird et al., 2017; Qi et al., 2017; An et al., 2019). Crops in the genus *Brassica* are also nutritionally and economically important worldwide, valuing around 14 billion dollars in 2014 (FAOSTAT 2014). One of the most widespread crop species, *Brassica rapa* L. (Brassicaceae: 2*n* = 20), includes turnips, leafy greens such as bok choy, napa cabbage, mizuna, tatsoi, rapini, grelos, and choy sum, and oilseed crops such as turnip rape, toria, and yellow sarsons as well as weedy forms (*B. rapa* ssp. *sylvestris*) which may be wild or feral (Table 1; Gómez-Campo and Prakash, 1999; Prakash et al. 2011). The diverse forms of *B. rapa* vary in both self-compatibility and annual/biennial habit. With the exception of the Central Asian oilseed crop, yellow sarsons, the species is largely reported as self-incompatible (Nasrallah and Nasrallah, 1989). Many of the crops and weedy forms are annual, but turnips, some turnip rape cultivars, and some napa cabbage are biennial and require vernalization to flower (Zhao et al. 2007). Propagation of *B. rapa* crops and weeds is through seeds and pollination of *B. rapa* is through insects and, to a lesser extent, wind and physical contact (Pertl et al. 2002). Despite the global economic importance of *B. rapa* as a crop species and its potential as a model system for artificial selection, many aspects of its domestication are contested. Outstanding questions include when, where, and how many times it was domesticated into various morphotypes, distinguishing the wild and feral types, what the first crop type was, how other crop types emerged from the initially-domesticated crop type, and to what degree different domesticated and feral forms have evolved convergently.

**Table 1.** Infraspecific taxonomy of *Brassica rapa* modified from Diederichsen (2001). Classification under the species level in *B. rapa* is currently contested (McAlvay et al., 2017), for example, the Iberian leafy crop known as *grelos* or *nabiza* are typically considered in the same subspecies as Italian *rapini*. See discussion for taxonomic implications of our results. *For simplicity brown sarsons and toria will be considered together as “toria.” ^†^For the purposes of this study, we define “weedy” *B. rapa* as wild or feral populations. We chose the term “weedy” over “wild” to include the possibility of ferality, and reflect the preference of spontaneously occurring *B. rapa* for disturbed areas (Weis and Kossler, 2004).

| Geography | Common name | Latin binomial | Plant organ(s) used |
| --- | --- | --- | --- |
| C. Asia | Yellow sarson | *ssp. trilocularis* (Roxb.) Hanelt | Seeds/leaves |
| C. Asia | Toria/Brown sarson* | *ssp. dichotoma* (Roxb.) Hanelt | Seeds/leaves |
| E. Asia | Komatsuna | *ssp. perviridis* Bailey | Leaves/inflorescences |
| E. Asia | Bok Choy | *ssp. chinensis* (L.) Hanelt | Leaves |
| E. Asia | Napa cabbage | *ssp. pekinensis* (Lour.) Hanelt | Leaves |
| E. Asia | Choy sum | *var. parachinensis* (L.H. Bailey) Hanelt | Leaves/inflorescences |
| E. Asia | Mizuna | *ssp. nipposinica* (L.H. Bailey) Hanelt | Leaves |
| E. Asia | Tatsoi | *ssp. narinosa* (L.H. Bailey) Hanelt | Leaves |
| E. Asia | Taichai | *ssp. chinensis* Makino *var. tai-tsai* Hort | Leaves |
| E. Asia | Zitaichai | *var. purpuraria* (L.H. Bailey) Kitam | Leaves/inflorescences |
| Europe | Rapini | *ssp. sylvestris* L. Janch. *var. esculenta* Hort | Leaves/inflorescences |
| Eurasia | Turnip rape | *ssp. oleifera* (DC.) Metzg. | Seeds |
| Eurasia | Turnip | *ssp. rapa* | Root-hypocotyl/leaves |
| Eurasia | Weedy (wild/feral)<sup>†</sup> | *ssp. sylvestris* (L.) Janch. | Leaves/inflorescences |

Past studies of *B. rapa* have recovered conflicting morphological, genetic, and geographic patterns. Whereas some studies show that morphologically similar crops (e.g., oilseeds) are closely related to each other despite being geographically distant (Zhao et al., 2005; Del Carpio et al. 2011; Tanhuanpää et al., 2015), others found that geographically proximal crops were closely related despite strong morphological differences (Song et al. 1988, 1990; Zhao et al. 2005; Takuno et al. 2007; Del Carpio et al. 2011; Annisa et al. 2013; Guo et al. 2014). This has led some researchers to propose multiple domestication events (Song et al., 1988; McGrath and Quiros 1992; Zhao et al., 2005), while others support a single domestication (Burkill and Haniff, 1930; Ignatov et al., 2008; Qi et al., 2017).

The proposed region(s) and timing of domestication are similarly varied. Eurasia harbors a number of centers of crop domestication including Southeast, East, South, and Central Asia, with the former two regions originating as early as the Early Holocene (Larson et al. 2014). Prior to the establishment of Silk Road networks linking all of these putative domestication centers, there was widespread early diffusion of crops across Europe and Asia. For example, certain Near-Eastern domesticates like barley are thought to have spread throughout Eurasia as early as 4000 years ago (Lister et al 2018). Proposed centers of origin for *B. rapa* include Europe (Song et al., 1988, 1990; Zhao et al., 2005), West Asia (Harberd, 1972), Central Asia (Ignatov et al., 2008; Qi et al., 2017), and East Asia (Song et al., 1988; Zhao et al., 2005). The timing of domestication has been difficult to assess due in part to the relative difficulty of finding and identifying *Brassica* seeds at archaeological sites (but see Allchin, 1969 and Hyams, 1971), but literary and linguistic evidence have provided some insights (Supp. table 1; Qi et al., 2017).

The initial crop type selected, its diffusion, and the subsequent selection for different crops is also debated. While some hypothesize that turnips were the first domesticated *B. rapa* crop type (McGrath and Quiros, 1990; Siemonsma and Piluek, 1993), early literary and linguistic evidence suggest that oilseed types are also ancient (Supp. table 1). Some genetic studies have variously suggested that napa cabbage was selected from bok choy (Song et al. 1990; Zhao et al. 2005; Takuno et al. 2007), turnips (McGrath and Quiros, 1990), or a cross between bok choy and turnips (Li, 1981; Song et al., 1988; Ren et al. 1995; Qi et al., 2017). Furthermore, several *B. rapa* leafy crops exist in northwestern Spain and Portugal (Francisco et al., 2009), but their origins have not been investigated1. Uncertainty also remains surrounding the *B. rapa* oilseed crops, as there are apparent references to yellow sarsons (*B. rapa* ssp. *trilocularis*) as far back as 2900 years ago in India, whereas turnip rape (*B. rapa* ssp. *oleifera*) may have originated relatively recently in Europe and China from turnips and leafy vegetables respectively (Gómez-Campo and Prakash, 1999). While an apparent multiple origin of turnip rape is widely supported by molecular data, the specific progenitors of the different lineages are unclear (Bird et al. 2017; Qi et al. 2017).

A major limitation to insights into the domestication of *B. rapa* has been a lack of knowledge about weedy forms (*B. rapa* ssp. *sylvestris*) which could be either truly wild or feral (McAlvay et al., 2017). Wild crop relatives can be key to clarifying the location and timing of domestication events (Meyer et al., 2012). The identification of teosinte as the progenitor of maize led to key insights into the morphological and genomic transitions of maize domestication and made substantial contributions to breeding efforts (Wilkes, 1967, Cruz-Cárdenas et al. 2019). Although *B. rapa* ssp. *sylvestris* occur spontaneously on roadsides, waste areas, farmlands, and riversides in many temperate areas worldwide (Andersen et al., 2009), it is unclear whether these populations are feral crop escapes or truly wild forms (Crouch et al., 1995; Andersen et al., 2009). Feral plants—derived fully or partially from domesticated crops—may or may not resemble wild forms and may harbor different levels of diversity than wild conspecifics (Gressel, 2005). Definitively distinguishing wild and feral populations can be challenging, but evidence for ferality may include crops emerging as sister to weedy populations in phylogenies or introgression between crop and weedy populations (Page et al. 2019). Disentangling feral and truly wild *B. rapa* could position the species as a powerful model system for studying both evolution under domestication and ferality given its close relationship to the genetic model *Arabidopsis thaliana*, rapid life cycle, and well-annotated reference genome.

To clarify the wild or feral status of weedy *B. rapa* populations and reconstruct the domestication history of crop forms, we performed genotyping-by-sequencing (GBS) (Elshire et al., 2011) on an unprecedentedly broad panel of *B. rapa* crops and weedy samples to assess genetic structure, diversity, introgression, and demographic history. We also modeled the distribution of weedy *B. rapa* in Eurasia and North Africa during recent and mid-Holocene conditions to project areas of suitable habitat during the putative time of domestication and identify potential modern distribution. We investigated the following questions: 1) Are weedy *B. rapa* populations wild or feral? 2) When, where, and how many times was *B. rapa* domesticated and what was or were the first domesticated crop type(s)? and 3) Were the crop morphotypes (leafy, turnip, oilseed) selected one or multiple times from the initial domesticated morphotype?

## Results

### Genotyping-by-sequencing

#### SNP calling, SNP filtering, and sample filtering

Sequencing produced 656,335,837 raw reads (average 218,778,612 per lane) that were combined with 823,954,356 raw reads from Bird et al. (2017). A total of 372,182 SNPs were called using the Tassel pipeline (Glaubitz et al., 2014). After downstream SNP and sample filtering, we retained 416 samples (Supp. table 2) and 68,468 SNPs at a minimum minor allele frequency of 0.05.

#### Diversity

Nucleotide diversity (Supp. table 3) within populations of *B. rapa* showed higher diversity in turnip samples from Central and Western Asia, intermediate levels of diversity in European crops and weeds, and lowest diversity in East Asian crops and South Asian oilseeds. The AMOVA analysis (Supp. table 4) revealed that most of the variation present is within populations (52.39%), followed by geographical groups (28.54%), and among populations within geographical groups (19.07%).

#### Genetic structure and maximum likelihood trees

To investigate the genetic structure of crop and weedy *B. rapa*, we carried out fastSTRUCTURE analysis across K values of 1 through 20. A K value of five maximized marginal likelihood in fastSTRUCTURE (Supp. fig. S1) with K = 8 best explaining the structure in the data using a variational inference algorithm. The resulting plots from K=5 to K=8 are shown in fig. 1. At K = 5, napa cabbage, bok choy, and yellow sarsons formed relatively discrete cluster, with a composite cluster consisting of toria and East Asian turnips/Japanese greens. Toria was unique from East Asian turnips/Japanese greens in that it has a small amount of shared ancestry with yellow sarsons, while East Asian turnips/Japanese greens share a small amount of ancestry with napa cabbage and bok choy. Central Asian turnips, weedy populations from Caucasus, Siberia, and Italy (together referred to as “weedy CSI” below), and Turkish turnips show shared ancestry with European crops and weeds, toria, the East Asian turnips/Japanese greens cluster, napa cabbage, and bok choy. At K = 6 differentiation emerges between different subclusters of napa cabbage with the purple cluster predominantly from South Korea and the red cluster largely from China and other countries. At K = 7, East Asian turnips and Japanese greens emerge in their own cluster. At K = 8, some of the Central Asian turnips (from Afghanistan) emerge as a separate cluster with some signatures of admixture with this Afghani cluster showing up in Turkish turnips, other Central Asian turnips, and weedy Caucasus, Siberia, and Italy.

**Figure 1.**
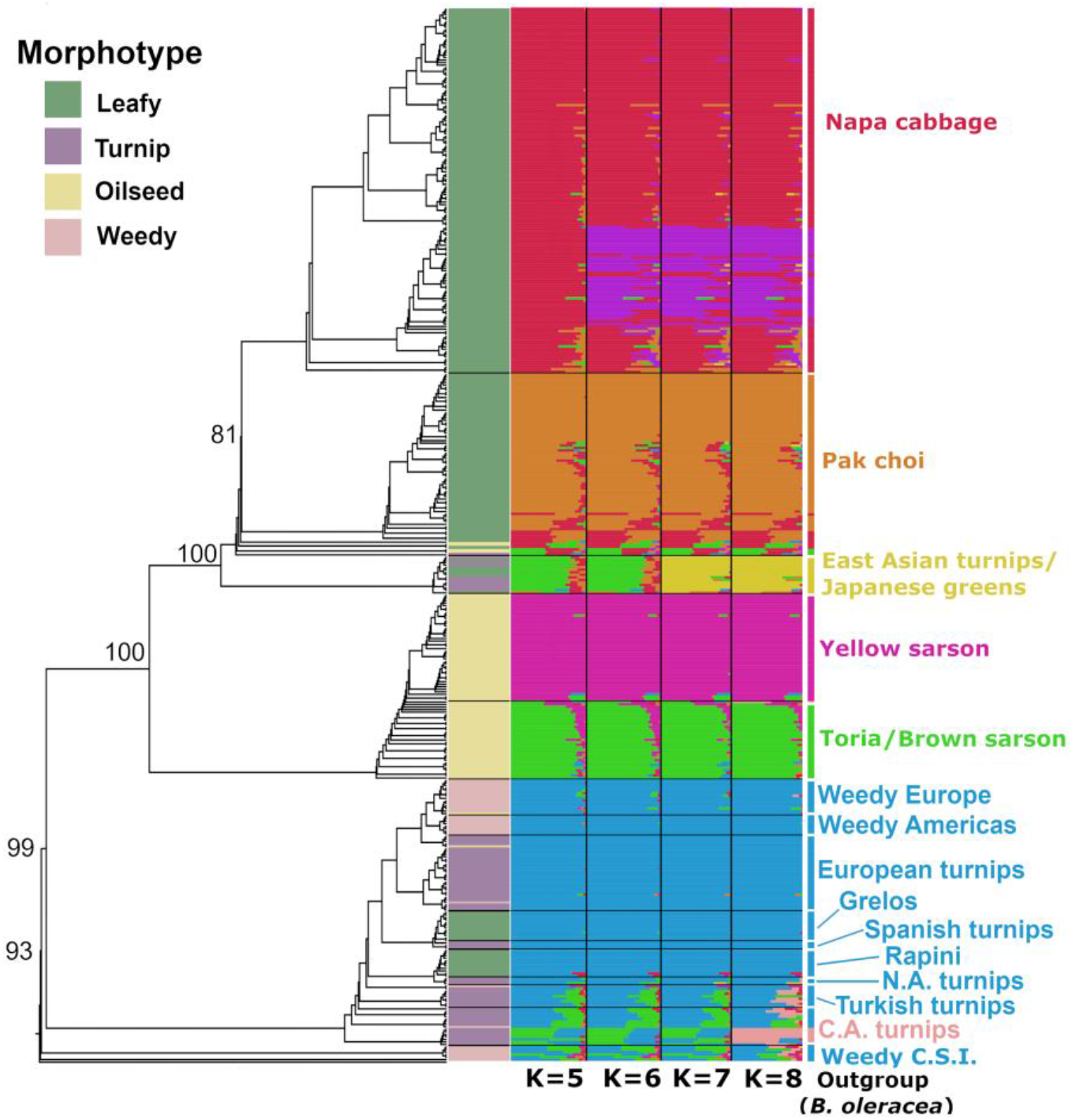
RAxML tree and fastSTRUCTURE plot indicating population structure of *Brassica rapa* crops and weeds at four different values of K. Each row represents a single individual and colors indicate its proportion of similarity with a given cluster. Bootstrap values for major clades are indicated as numbers at nodes. C.A. turnips indicates Central Asian turnips and weedy C.S.I. indicates weedy samples from the Caucasus, Siberia, and Italy. Turnip rape (*B. rapa* ssp. *oleifera*) samples are not indicated in the labels at right as they emerged sporadically in small numbers in several parts of the tree (with European turnips, weedy Europe, and bok choy). These samples can be seen in the figure as the oilseed samples (as indicated in yellow in the morphotype bar) that fall outside of the toria/yellow sarson clade. Weedy C.S.I., weedy Europe, and weedy Americas are all *Brassica rapa* ssp. *sylvestris*.

To investigate the relationships between individuals, we built maximum likelihood trees in RAxML and SNPhylo as well as a distance-based neighbor-joining (NJ) tree. In the RAxML tree (fig. 1), the weedy CSI samples are recovered as sister to the remaining lineages with strong bootstrap support (BS; 93%). Weedy European and American populations did not cluster with the weedy populations from CSI, instead emerging within the European crop clade. European turnips were sister to the weedy European and American populations with low BS (35%). Central Asian turnips emerged as sister to a clade of North African and Turkish turnips, European crops and European weedy populations. The leafy European crops (rapini and grelos) emerged in separate clades with North African turnips sister to rapini (97% BS) and Spanish turnips sister to grelos (56% BS). Toria emerged in a grade leading up to a clade of self-compatible yellow sarsons with short branches. The yellow sarson and toria were together sister to the East Asian crops. The East Asian Turnips and Japanese greens clade was sister to clade containing napa cabbage and bok choy (100% BS). The SNPhylo tree (Supp. fig. S2) was broadly congruent with the RAxML tree except weedy CSI, Central Asian turnips and Turkish turnips emerged as sister to East and Central Asian crops rather than with weedy CSI sister to all crops and weeds and Central Asian and Turkish turnips as sister to all European crops and weeds. The NJ tree (Supp. fig. S3) largely corroborated the RAxML results with minor differences. The principal difference was that in the NJ tree, North African turnips came out as sister to weeds from the Americas and Europe rather than to rapini.

The PCA generally separated samples based on geography and crop type (fig. 2). PC1 (18.3% of variance explained) separated Central Asian oilseeds from the remaining crops and weeds. PC2 (10.4% of variance explained) separated East Asian crops from Central Asian turnips, European turnips, greens, and weeds. The weedy CSI samples clustered in between Central Asian turnips, toria, and East Asian turnips/ Japanese greens, while weedy forms from Europe and the Americas were closely affiliated with European turnips and Mediterranean leafy types (Italian rapini and Spanish grelos). The European weeds and turnips are not well differentiated from each other using the first several PC axes (Supp. fig. S4A). Among East Asian crops, turnips emerged as most closely associated with Central Asian crops and weeds, followed by bok choy, and napa cabbage. Subsets of the samples are shown in Supp. fig. S4B-F.

**Figure 2.**
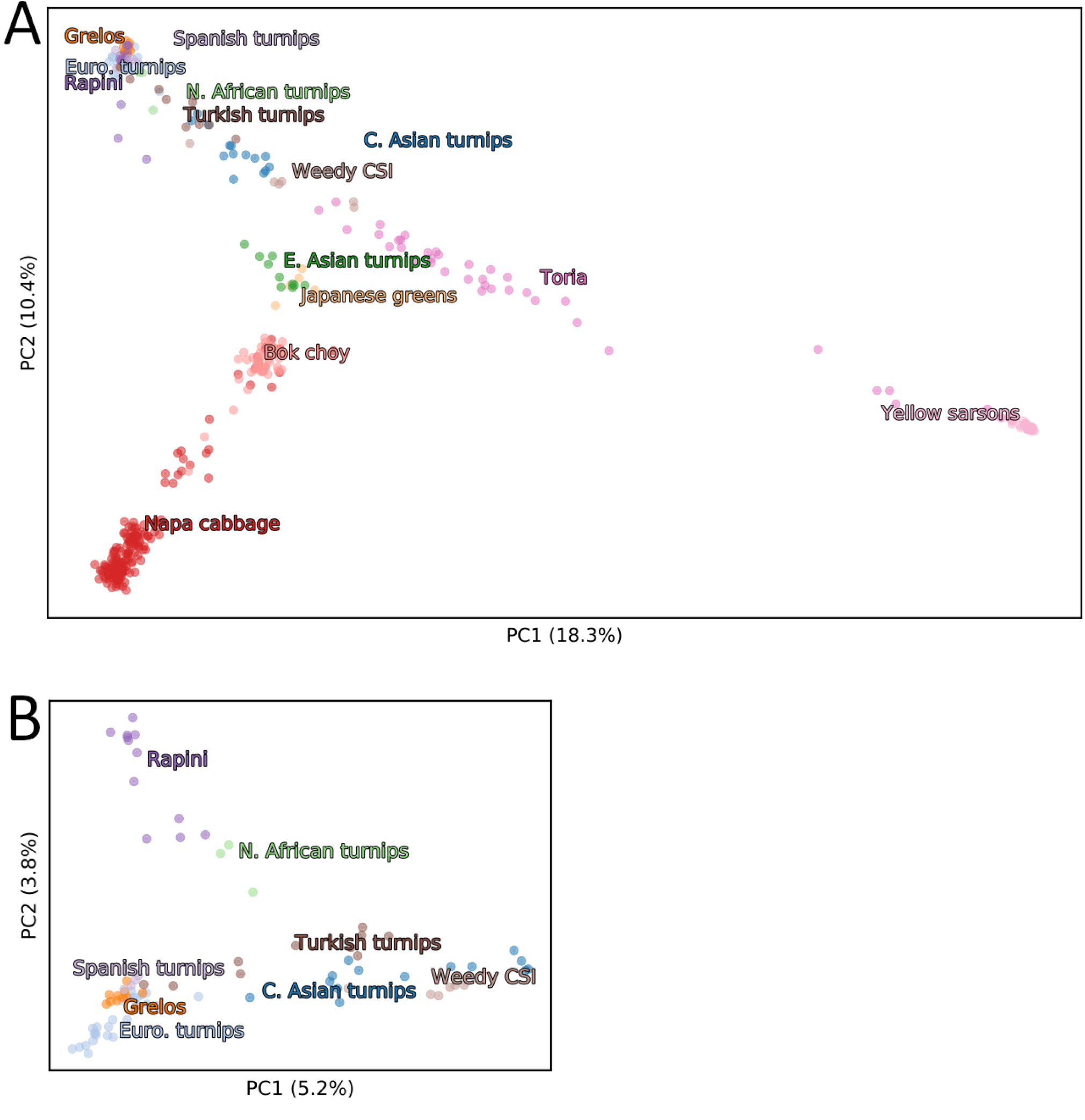
(A) Principal component analysis displaying PCs 1 and 2 of *Brassica rapa* samples using global crops and weedy forms (B) and focusing on Central Asian and European crops. When considering all populations, PC1 separates sarsons and toria from other crops and feral forms, and PC2 separates the Eastern and Western regions. When focusing on just Central Asian and European turnip and leafy crops (B), PC1 captures the geographic cline from West to East, while PC2 separates rapini and North African turnips from grelos and other European turnips. Additional plotted PCs are shown in Supp. fig. S4. The color of labels corresponds to the colors of the points of associated accessions.

F_ST_ values were significantly different among populations and ranged from 0.046 between European weeds and European turnips to 0.899 between yellow sarsons and North African turnips (Supp. table S5). Grelos were least differentiated from Spanish turnips and rapini were least differentiated from other European turnips. Yellow sarsons were largely differentiated from all other morphotypes (all values >0.5). Weeds from the Americas showed lowest differentiation from Spanish turnips (0.076) and European weeds (0.071). Weedy populations from CSI were least differentiated from Central Asian turnips (0.126), European turnips (0.118), and East Asian turnips (0.119).

#### TreeMix and f_4_-statistics

To investigate reticulation in the evolutionary history of *B. rapa* crops and weeds, we used TreeMix (Pickrell and Pritchard 2012) and four-population (*f*_*4*_) tests of treeness (Reich et al. 2009). TreeMix generates phylogenetic networks by first creating a dendrogram based on genetic drift and then comparing the covariance structure of the dendrogram to covariance across populations to estimate admixture. *f*_4_-statistics test the support for admixture in tested populations by detecting whether allele frequency correlations are compatible with treelike evolution. Overall, the relationships recovered are largely congruent with the RAxML phylogeny of Eastern and Western clades. However, the major difference is the placement of weedy CSI as sister to the Asian or Eastern clade rather than as sister to all other cultivars as in the RAxML (fig. 1) and NJ tree (Supp. fig. S3). The tree model with no migration explains 96.4% of the variation in relatedness between the populations, however sequentially adding migration events results in four migration events representing 98.5% the variation (Supp. fig. S5). Two migration events are supported by four-population (*f*_4_) tests for treeness (fig. 3): between toria and Central Asian turnips (*f*_*4*_= -0.003, Z = -8.03) and between weedy CSI and weedy European populations (*f*_*4*_= -0.003, Z = -5.98)

**Figure 3.**
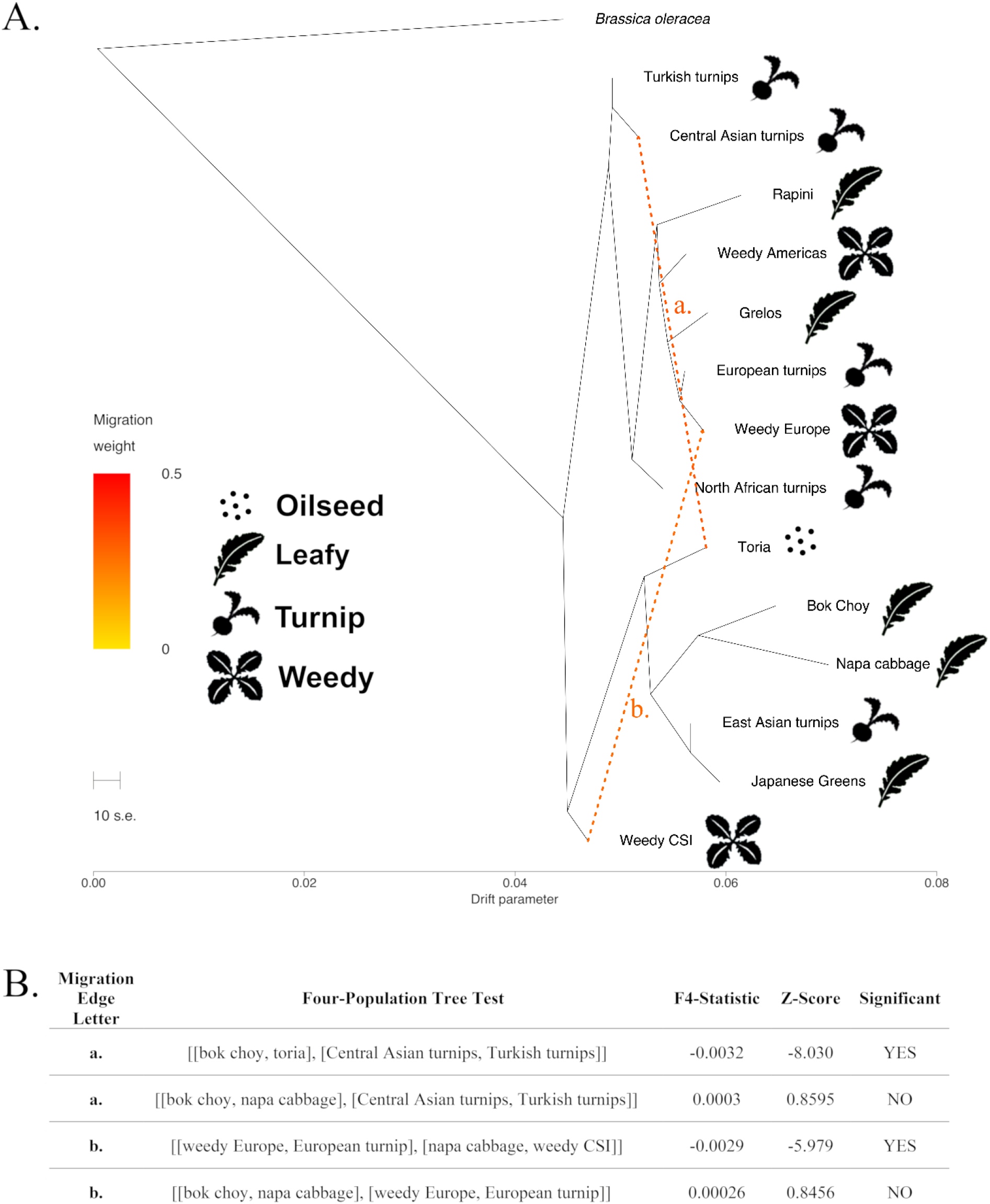
TreeMix diagram and *f*_*4*_ -statistics. A) TreeMix diagram indicating relationships between *B. rapa* crop and weedy forms with recovered migration events indicated with orange dotted lines. B) *f*_*4*_ -statistic results for migration edges recovered in TreeMix.

#### Linkage disequilibrium decay, effective population sizes, and demographic modeling

In order to investigate population branching structure, population size changes, and migration between populations, we constructed and tested demographic models using *moments* v1.0.1 (Jouganous et al. 2017), which uses the joint allele frequency spectrum (AFS) to fit parameterized demographic models for the populations depicted in Supp. table S9. To estimate *Ne* and visualize patterns of linkage disequilibrium (LD), we also calculated and graphed linkage disequilibrium for populations. Our LD analysis revealed that yellow sarsons have strongly elevated long-range LD, while toria and East Asian turnips also show elevated long-range LD in comparison to other populations (Supp. fig. S6). Using LD between unlinked SNPs, those populations also were also found to have the smallest recent population sizes with *Ne≅*5-10 (Supp. table S6). Yellow sarsons are self-compatible (Sethi et al., 1970), which can dramatically increase long-range LD and result in reduced estimates of *Ne* (Ragsdale & Gravel, 2020), while high levels of inbreeding or a recent strong bottleneck may also increase long-range LD and lead to small estimates of *Ne*.

For the demographic models comparing toria, turnips, and weedy CSI, we found that models that allowed for migration always fit far better than those that did not allow for migration, that the inferred migration rates are very large, and in general we see that the admixture models for weedy CSI are not better supported than divergence with gene flow. However, we cannot confidently infer the relative order of splits between toria, weedy CSI, and turnip crops, likely because the very high migration rates that extend from the time of divergence to the present obscure any signal of a “clean” split between populations (Supp. table S7).

We inferred demographic models for sets of two to four populations using the joint SFS with *moments* (Jouganous et al. 2017). We first focused on populations in central Eurasia to try to disentangle the deeper population structure and relative order of divergences between toria, turnips, and weedy CSI (Supp. table S7). In general, our best fit models show that there were high levels of gene flow between toria, turnips, and weedy CSI, with only European crops having more moderate gene flow with Central and East Asian populations. Because of the large gene flow between toria, turnips, and weedy populations, we were unable to definitively resolve the relative timing of splits between these populations. Our best fit models do not support the scenario that weedy CSI arose recently through admixture, but instead that its existence predates the more recent radiation of turnip and leafy crops in Europe and East Asia.

We then turned to inferring demographic histories of turnip and leafy crops (Supp. table S7, Supp. fig. S7). Across all tested demographic models, we inferred that crop forms split from weedy *B. rapa* from the Caucasus, Italy, and Siberia around 3,430-5930 years before present, assuming on average one generation per year. The European and Asian lineages diverged roughly 1,930-2,430 years ago, with subsequent diversification over the last one to two thousand years. However, there is large uncertainty surrounding these inferred dates. Not only are the per-base mutation rate and long-term generation times uncertain, the very high levels of inferred migration confound the inferred split times.

For the more recent history, we observe multiple origins of leafy green crops in both the Western and Eastern regions. Grelos split from Spanish turnips roughly 730-930 years before present, with subsequent high levels of gene flow between those two lineages, while rapini split from the Turkish turnip lineage around the same time, with subsequent gene flow between rapini and European turnips. In the Eastern lineage, we observe an early diversification of napa cabbage from turnips, and a more recent emergence of bok choy, with contributions from both East Asian turnips and napa cabbage.

We inferred high levels of regional migration between nearby crops and reduced or zero recent gene flow between more distant lineages. For example, we inferred high levels of gene flow between Spanish and European turnips, and between European and Turkish turnips, but far less direct gene flow between Spanish turnips and turnips from Turkey or farther east. Similarly, we inferred migration between East and Central Asian turnips, but not between East Asian turnips and crops in Europe.

#### Species distribution modeling

Species distribution modeling or environmental niche modeling uses species occurrence data based on herbarium specimens, observations, and other occurrence data combined with environmental data to predict suitable habitat in the past, present, and/or future (Kozak et al., 2008, Nakazato et al., 2010). In order to narrow the potential center of domestication and identify conservation priority areas for wild relatives, we generated species distribution models to infer the fundamental ecological niche of *B. rapa* under current and mid-Holocene climate conditions. The omission rate on training samples was close to predicted omission and the receiver operating characteristic curve (DeLong et al., 1988) showed an area under curve of 0.957, suggesting that the model fit the data well (values above 0.75 are considered informative) (Phillips and Dudik, 2008). The following three bioclimatic variables were most important for describing the niche: mean temperature of driest quarter (28.7%), mean temperature of coldest quarter (26.7%), precipitation of coldest quarter (10.8%. The model predicting *B. rapa* distribution under a mid-Holocene climate scenario (fig. 4A) indicated suitable habitat in a nearly contiguous band from Iberia, North Africa, and the British Isles to coastal China and Japan. This strip of habitat mainly followed highland formations, except in Europe, where coastal areas were also of high suitability. A potential gap in suitable habitat was present in the present-day borderlands of Afghanistan and Iran. The contemporary model (fig. 4B) was generally similar to the mid-Holocene model except for a general pattern of greater suitability in northern areas and lesser suitability in southern areas.

**Figure 4.**
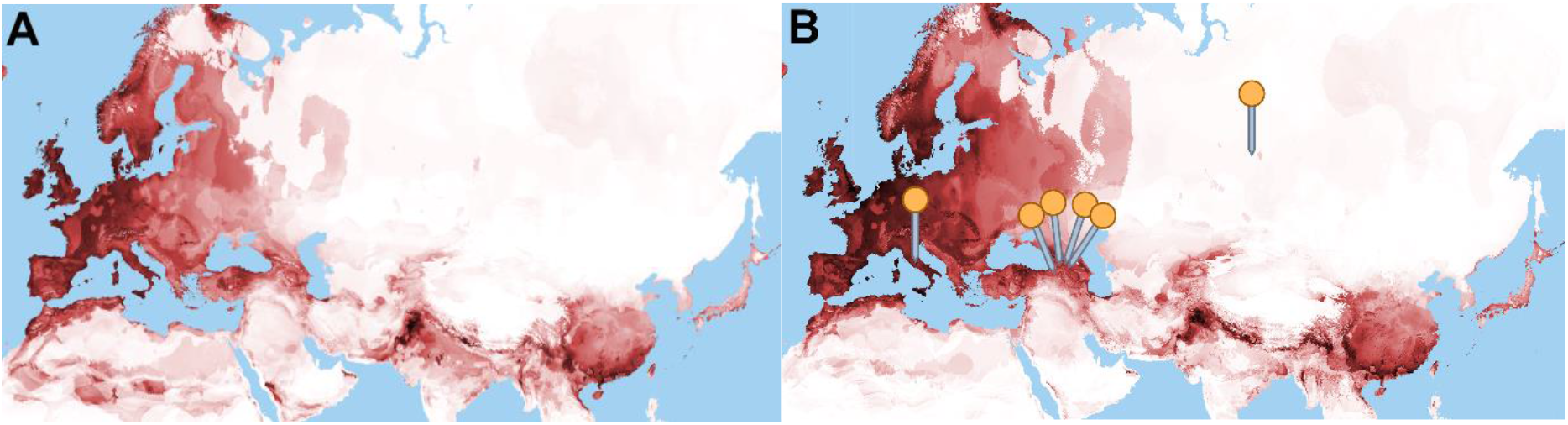
Environmental niche model for *Brassica rapa* in Eurasia and northern Africa for mid-Holocene (∼6000ybp) climate conditions (A) and contemporary climate conditions (B). Darker areas indicated better niche fit for weedy *B. rapa* with black indicating highest habitat suitability and white indicating poorest habitat suitability. Orange pins indicate areas where individuals from the “Weedy CSI” clade were collected.

## Discussion

### Insights into the identities of wild and feral populations

The identification of crop wild relatives allows for direct comparisons between wild and crop forms and provides an important source of genetic material for crops (Honnay et al., 2012; Dempewolf et al., 2014) which often have relatively limited genepools due to diversity bottlenecks from selection (Olsen and Gross 2008). Feral relatives can be morphologically indistinguishable from wild material (Wang et al. 2017) and the wild or feral nature of *B. rapa* populations have been contested (Andersen et al., 2009) hindering domestication research, breeding, and conservation in the species. In our analyses, weedy *B. rapa* samples from the Caucasus, Siberia, and Italy (“weedy CSI”) consistently segregate in a lineage distinct from weedy samples from the Americas and Europe outside of Italy, a lineage that has not been recovered in previous studies. With the exception of a single accession from Abruzzo in central Italy, and an accession from Tomsk in south-central Siberia, the weedy CSI clade consists of samples in or close to the Caucasus mountains in Georgia and northeastern Turkey. Both mid-Holocene and contemporary niche models suggested high habitat suitability in the Caucasus and Italy, but generally low suitability across Siberia except for in the extreme southern region. The lack of suitability in the latter case could be due to a lack of occurrence data from Siberia used in the niche model or erroneous passport data for the Siberian accession. Our analyses present the possibility that this lineage represents either truly wild relatives of some or all *B. rapa* crops or populations deriving from an early feralization event.

The hypothesis that the weedy CSI clade represents truly wild populations is consistent with Sinskaya’s (1969) suggestion that wild forms still exist in the Caucasus mountains and Siberia and our niche models indicate that habitat for *B. rapa* in the Caucasus, Siberia, and Italy would be suitable under a mid-Holocene climate model (fig. 4A). The position of weedy CSI samples as sister to all crops and weeds in the RAxML and NJ analyses, and high nucleotide diversity support this interpretation. This lineage is also located at the juncture of all the crop subspecies in the PCA and contains ancestry common to each of the major crop clades in fastSTRUCTURE, which may indicate that the other crops as weeds were subsampled from the diversity present in weedy populations in the CSI range. The placement of weedy CSI in the SNPhylo and TreeMix trees as sister to Central and East Asian crops, suggests the possibility that weedy CSI is truly wild, but an independent domestication event gave rise to the European crops.

An alternative interpretation would be that weedy CSI represents a highly admixed feral lineage. Many crops are known to exist in wild-domesticated-feral hybrid swarms (e.g., Beebe et al., 1997; Wang et al. 2017; Allaby et al. 2008). As mentioned above, the diverse ancestry for these individuals recovered by fastSTRUCTURE and the central position in the PCA could be explained by admixture between crop and/or weedy types. Several patterns traditionally associated with wild progenitors in analyses may instead indicate admixed feral populations (Wang et al. 2017). For example, high levels of admixture could also have inflated nucleotide diversity and caused the clade to be placed at an artificially deep position in phylogenetic analyses. Demographic models tested in *moments* were inconclusive regarding the early splits of the major lineages, but the models that performed best allowed for extensive early gene flow, supporting this hypothesis. The TreeMix recovered a migration edges supported by *f*_*4*_-statistics that would suggest gene flow between CSI and toria, and the TreeMix and SNPhylo trees are consistent with weedy CSI populations deriving from an ancient feral lineage that branched off from an ancestor of Central Asian oilseeds and East Asian crops.

Our analyses recovered strong evidence for the feral nature of the weedy populations outside of the CSI clade. Non-CSI European and American weedy samples emerged in association with European crops in PCA, fastSTRUCTURE, RAxML (99% BS), SNPhylo, and NJ analyses, and had relatively low differentiation from European crops in the F_ST_ analysis. Despite lower levels of diversity among the feral samples, their local adaptation to biotic and abiotic stresses could make them a valuable source of breeding material as in feral rice (Li et al., 2017).

Due to the geographic disjuncture of the weedy CSI samples, it is likely that there are related populations in the intervening areas. Several possibilities could explain the geographic disjuncture of the samples from Italy and the Caucasus in the CSI clade. Weedy *B. rapa* from the intervening area (e.g., Slovakia, Austria, and Serbia) were associated with other European weeds and crops in the analyses, but the TreeMix analysis suggested that introgression had occurred between the CSI clade and the European weeds, consistent with an exoferal origin of most of the European weedy accessions. Truly wild populations could have been more widespread in Europe previously, but may have been largely replaced by successful feral populations that are preadapted to agroecosystems. Alternatively, the sole Italian sample could have been introduced relatively recently as a seed contaminant in imported crops. Weedy populations of *B. rapa* have recently been reported in Southwest China (Dong et al. 2018) and Japan (Aono, 2011), despite previous reports of absence from East Asia (De Candolle 1886; McGrath and Quiros, 1992) as well as in Algeria (Aissiou et al. 2018). Sampling these populations could further clarify the evolutionary history of *B. rapa*.

### Domestication center, timing, and initial crop type

The location and timing of domestication as well as the initial domesticated form of *B. rapa* that was subsequently selected for other crop types has been debated. Our analyses support an initial domestication of turnips and/or oilseeds in Central Asia 3,430-5,930 YBP with subsequent diffusion to Europe and East Asia.

Turnips from Central Asia appear to be most closely associated with the putatively wild CSI populations in our F_ST_ analysis and had the highest levels of nucleotide diversity of any population. This is consistent with the findings of Qi et al. (2017), who found Central Asian and European turnips to have relatively higher values than East Asian vegetables, and much higher values than yellow sarsons and toria. Given our use of different reduced representation markers, the absolute comparison of values is not possible. Also as in Qi et al. (2017), we estimated highest Ne for turnips, intermediate values for East Asian vegetables, and the lowest values for yellow sarsons and toria. The turnip samples from the Hindu Kush mountains of Afghanistan, Pakistan, and Tajikistan are sister to the rest of the *B. rapa* samples from West Asia and Europe in our TreeMix, RAxML, and NJ trees and demographic model and are located in an area of very high habitat suitability in the Mid-Holocene species distribution model.

East Asian turnips are also associated with the putatively wild samples and Central Asian turnips as seen in the PCA, fastSTRUCTURE, and F_ST_ analyses. East Asian turnips also emerge as sister to other East Asian crops (with the exception of the Japanese leafy crops which are intermingled with East Asian turnips) in our tree-based analyses. An early Central Asian origin is supported by linguistic evidence for turnips in the Pontic Steppe north of the Caucasus mountains approximately between 6430-4278 YBP and in S.W. Asia nearly 3000 YBP (Supp. table 1).

It has been hypothesized that Central Asian oilseeds arose from an independent domestication event (Zhao et al., 2005; Warwick et al., 2008) or from European oilseeds (Song et al. 1988). While our TreeMix, SNPhylo, NJ, and RAxML trees show Central Asian oilseeds from the mountains of Afghanistan, Pakistan, and India as sister to East Asian crops, they are associated with West, Central, West, and East Asian turnips in fastSTRUCTURE. These results support either an independent domestication for oilseeds or selection on Central Asian turnips before the diversification of other crop forms across Europe and Asia. Literary evidence for oilseed crops in Northern India almost 3000 YBP supports their antiquity in the area (Supp. table 1). Apparent introgression between the Central Asian oilseed toria and Central Asian turnips as recovered by TreeMix and *f*_*4*_ statistics is consistent with the geographic proximity of these crops. In our tree-based analyses, toria is consistently sister to yellow sarsons.

In the demographic models we tested with weedy CSI as sister to the remaining crops and weeds, the split was estimated at between 3,430-5,930 YBP (years before present). If the weedy CSI populations are truly wild, this provides a rough temporal estimate of the domestication of *B. rapa*. While this prediction is sensitive to our estimates of initial effective population size and generation time, this finding is consistent with archaeological and linguistic evidence (Supp. table 1).

### Convergent phenotypes in leafy and oilseed crops deriving from turnip crops

Our inclusion of diverse leafy crop types from Europe and Asia allowed us to identify distinct evolutionary lineages for at least three different types of leafy *B. rapa* crops—grelos, rapini, and East Asian greens—all with likely origins in different turnip populations. None of our tree-based analyses recovered these three leafy crops together as a common monophyletic clade and they do not cluster with one another in the PCA. Instead, turnip crops from geographically similar regions emerged as sister to each one.

North African turnips emerged as sister to the Italian leafy crop rapini in TreeMix, RAxML (97% BS), SNPhylo, NJ analyses and Turkish turnips were sister to rapini in the *moments* demographic model. Our findings differ from Qi et al. 2017 who found rapini to be sister to the Central Asian oilseeds possibly because the majority of the rapini samples used in Qi et al. (2017) were later suspected to be polyploids in Bird et al. (2017) or had Central Asian provenance. Our findings suggest a possible trans-Mediterranean introduction of turnips to Italy followed by selection for leafy forms or domestication for a leafy type in North Africa where turnip leaves are also eaten (Hammer and Perrino 1985). A different Mediterranean turnip population, Spanish turnips, however, emerged as sister to the Spanish leafy crop grelos in our RAxML, SNPhylo, and NJ analysis. PCA results also indicate an association of the two Mediterranean leafy crops with distinct turnip populations. This provides the first evidence that grelos were independently selected from local turnip crops instead of sharing a common origin with rapini (Francisco et al., 2009).

A similar pattern was recovered in East Asia, where leafy crops may have an origin in a distinct population of turnips. East Asian turnips emerged as sister to East Asian leafy crops in the TreeMix, RAxML, SNPhylo, and NJ trees consistent with the results from Bird et al. (2017) and Cheng et al. (2016). The origin of East Asian leafy types in turnips is also consistent with Takuno et al.’s (2007) hypothesis that an unimproved proto-crop from Europe or Central Asia was introduced to East Asia where it was subsequently selected into diverse leafy morphotypes. The origins of napa cabbage as a result of intermixing of bok choy and East Asian turnips is supported in our demographic model as in Qi et al. (2017), but not recovered in the TreeMix migration edges supported by four-population tests. Most Japanese leafy greens are also clustered with the East Asian turnips suggesting that they too may share a common origin in turnips and may have been selected from an early introduction of turnips or a proto-vegetable type. While the East Asian turnips sampled had elevated LD compared to bok choy and napa cabbage, Cheng et al. (2016) found the opposite, perhaps as a result of their broader sampling.

While these patterns could suggest parallel selection for leafy types out of distinct turnip populations, they may also be explained by a single origin of the leafy morphotype with subsequent introgression resulting in the leafy phenotype arising in other turnip lineages, although the latter possibility is not supported by our TreeMix and *f*_*4*_-statistic analyses. The enlarged root-hypocotyl of turnips is controlled by relatively few genes and may have therefore been lost multiple times through human selection and/or ferality (McGrath and Quiros 1991). The pattern of selection for leafy crops out of conspecific crops with swollen stems or taproots does not seem to be common in analogous crops. A similar trend has been observed, however, in *B. juncea* as it appears that root mustard (*B. juncea* ssp. *napiformis*) was the initially selected domesticated, followed by leafy, stem, and oilseed types (Yang et al. 2018). This differs from *B. oleracea, B. napus*, and *Beta vulgaris* where leafy or oilseed types appear to predate the swollen stem/root types (Zohary and Hopf, 2000; Maggioni, 2015; An et al., 2019).

Our limited sampling of turnip rape oilseeds from East Asia and Europe restrict our inferences, but like Bird et al. (2017) and Qi et al. (2017), turnip rape did not form a monophyletic clade. Our data supports an origin of European turnip rape from European turnips, and East Asian turnip rape from turnips and/or bok choy consistent with Reiner et al. (1995).

### Taxonomic implications

Our findings indicate that an infraspecific taxonomic recircumscription of *B. rapa* is warranted to avoid confusion. Several classification systems are currently in use with a combination of subspecies, varieties, and cultivar groups as infraspecific designations (McAlvay et al. 2017). Our findings highlight several paraphyletic lineages that are typically considered to be the same taxon including turnip rape (*B. rapa* ssp. *oleifera*), weedy forms (*B. rapa* ssp. *sylvestris*), and rapini/grelos (*B. rapa* ssp. *sylvestris* var. *esculenta*). In the case of the latter name the ‘ssp. *sylvestris*’ epithet is misleading given its likely (multiple) origins in turnips. Furthermore, our results indicate that turnips may also have multiple origins or a single origin with reversions to turnip types, with the former scenario suggesting the need for a reexamination of nomenclature. A phylogenetically and morphologically informed recircumscription of *B. rapa* to create an international standard for delineating these lineages would be a valuable future contribution to *Brassica* research and breeding.

## Conclusions

Our broad sampling of crops and weedy populations of *Brassica rapa* allowed us to identify feral and potentially wild relatives, identify a putative center of domestication, and reconstruct the diversification history of crop morphotypes. These insights enhance *Brassica rapa* as a model system to study artificial selection and ferality as well as contribute to conservation of crop genetic resources in this species.

We found evidence for a distinct lineage of *B. rapa* from the Caucasus, Siberia, and Italy which may represent the first documented wild *B. rapa* populations or the descendants of an ancient feral lineage, and we determined that weedy plants in a large portion of the European and American distribution are feral populations resulting from introgression between relatives of weedy CSI and European turnips. Our findings provide evidence of the complex interplay of wild, crop, and feral populations that may take place throughout the domestication process and raise the possibility that the evolutionary relationships between these lineages is not treelike, a pattern also observed rice (Wang et al. 2017), common bean (Beebe et al., 1997), and other crops. If the weedy CSI lineage is truly wild, it provides an unprecedented opportunity to directly compare wild, domesticated, and feral plants to determine functional genomic differences and morphological changes under human selection and feralization (Page et al., 2019). This enhances *B. rapa*, which has a well-characterized genome, short generation time, and diversity of crop types, as a powerful model system for understanding domestication and ferality. Distinguishing these gene pools contributes to the prioritization of *B. rapa* populations for conservation and germplasm collection (Brozynska et al, 2016) that may prove important in the future to cope with changing environmental conditions (Guarino and Lobell 2011). This is especially urgent, as potentially wild populations of *B. rapa* that are seen as weeds could be harmed by increased agricultural weed control in its native range (Crouch et al. 1995).

We found evidence for a common origin of *B. rapa* crops in turnips or oilseeds domesticated in the mountains of Central Asia between 3,430 and 5,930 years ago with subsequent selection on turnips for leafy vegetables and oilseeds one or multiple times in the Mediterranean and East Asia. Our findings highlight the importance of Central Asia for germplasm collection and in situ agrobiodiversity conservation and indicate, for the first time, that the Spanish leafy crop grelos has a distinct origin from Italian rapini. The appearance of various geographically disparate but morphologically convergent leafy, oilseed, and tuber-forming types suggests that crops in other species with multiple morphotypes such as other *Brassica* species and *Beta vulgaris* (beets, chard), may have been selected for particular suites of traits on multiple occasions. The nutritional characteristics, culinary uses, and storability of these organs in each of these species is relatively different, suggesting that selection for multiple morphotypes may have been a common strategy to fill different food systems niches across Eurasia.

Our results raise new questions about the nature of the weedy CSI lineage, potential parallel selection, as well as domestication and feralization syndromes in *B. rapa*. Additional sampling of *B. rapa* populations in the Caucasus, Siberia, and Italy as well as known weedy populations in western China (Dong et al., 2018), Japan (Aono, 2011), and North Africa (Aissiou et al. 2018) would provide further clarity regarding wild and feral gene pools. Our finding of convergent leafy and oilseed phenotypes from distinct turnip lineages may suggest that parallel selection may have occurred, a hypothesis that could be tested explicitly with functional genomic approaches. Finally, our study opens up the possibility of understanding the genetic and phenotypic basis of domestication and feralization in *B. rapa* with direct comparisons possible between weedy CSI, turnip, and feral populations.

## Methods

### Genetic analyses

#### Sampling

We obtained 289 *B. rapa* and outgroup samples through seed banks (e.g., USDA, IPK-Gatersleben), and directly from researchers (Supp. material 1). Three seeds from each accession as well as three seeds from each of the 333 USDA seedbank accessions used in the final GBS dataset by Bird et al. (2017) were planted in 6” square pots using Promix HP medium (Premier Tech, Rivière-du-Loup, Québec) at the Walnut Street Greenhouses at University of Wisconsin-Madison. Accessions were characterized morphologically using select standard descriptors developed by the International Board for Plant Genetic Resources (IBPGR, 1980). Subspecies identity was evaluated (Supp. table S2) using the following criteria: semi-heading and heading with wrinkled leaves and swollen petioles—napa cabbage (*B. rapa* ssp. *pekinensis*), swollen petioles and smooth spatulate leaves—bok choy (*B. rapa* ssp. *chinensis*), parted leaf dissection—mizuna (*B. rapa* ssp. *nipposinica*), flattened rosette of many smooth spatulate leaves (*B. rapa* ssp. *narinosa*), long swollen petioles with smooth orbicular leaves—komatsuna (*Brassica rapa* ssp. *perviridis*), enlarged root-hypocotyl and dissected lyrate leaves—turnip (*B. rapa* ssp. *rapa*), compact and enlarged inflorescence with dissected lyrate leaves—rapini and grelos (*B. rapa* ssp. *sylvestris* var. *esculenta*), yellow seeds—yellow sarsons (*B. rapa* ssp. *trilocularis*), lacking the traits previously mentioned but possessing dissected lyrate leaves—toria or brown sarson (*B. rapa* ssp. *dichotoma*), turnip rape (*B. rapa* ssp. *oleifera*) or weedy (*B. rapa* ssp. *sylvestris*) based on collection passport data (e.g. country, cultivated/uncultivated context).

GBS data from these samples were combined with a previously generated GBS diversity dataset (Bird et al., 2017), providing a total of 653 samples before sample filtering. This diversity panel incorporates samples from six continents and 12 crop subspecies. Our panel augments Bird et al.’s (2017) sampling of East Asian leafy crops, South and Central Asian oilseeds, and European turnips by adding turnips from West Asia, South Central Asia (“Central Asia,” defined as Nepal, Northern India, Pakistan, and Afghanistan, Tajikistan, Kyrgyzstan, and Western China), and Europe, Mediterranean leafy crops (Spanish grelos and Italian rapini), and weedy occurring samples from Europe, the Caucasus, Siberia, the Americas, and New Zealand. We also included a *Brassica oleracea* sample (NGB 162411) from Denmark to serve as an outgroup for our tree-based analyses. We did not include three East Asian vegetable crops (caixin, zicaitai, taicai), which are thought to be selected from bok choy (Cheng et al. 2016). Choy sum also appears to be selected from bok choy and as a result is included with bok choy in our analyses.

#### DNA extraction and sequencing

DNA was extracted from young leaf material using the CTAB method (Doyle and Doyle, 1987) at the University of Wisconsin-Madison Biotechnology Center (UWBC). The restriction enzyme ApeKI was used to construct GBS libraries. Samples were 96-plexed in each of three lanes of Illumina HiSeq 2000 (Illumina Inc. San Diego, CA, United States), and one well in each plate was left as a negative control. Raw sequence data was combined with that of Bird et al. (2017), who also used the ApeKI enzyme, and processed using the GBS 2 pipeline in Tassel 5 (Glaubitz et al., 2014). Parameters used for the pipeline can be found in Table S8. Reads were aligned to *B. rapa* ssp. *pekinensis* v1.5 reference genome (Wang et al., 2011) using Burrows-Wheeler Alignment (Li and Durbin, 2009). Raw reads are publicly available on Data Dryad at https://doi.org/10.5061/dryad.0gb5mkm0m.

#### SNP and sample filtering

The SNPs resulting from the Tassel 5 pipeline were filtered using VCFtools (Danecek et al., 2011) based on read depth (minimum mean depth = 3), number of alleles (only biallelic loci used), minimum number of genotypes scored per site (100% for PCA, 90% for all other analyses), and a minimum minor allele frequency of 5% for all analyses except demographic modeling (Supp. table S8). In order to test the effects of different SNP filtering criteria, we ran the PCA, RAxML, and FastSTRUCTURE analyses with SNP datasets that had been filtered with minimum mean read depths of 3, 6, 9, and 12. We found minimal differences in the patterns recovered by the three analyses and so retained the minimum mean read depth of 3. Tassel 5 was used to filter SNP sites for a maximum heterozygosity of 50% and remove taxa that had <50% of scored loci. Samples identified by Bird et al. (2017) to be polyploids or otherwise problematic were removed (see Bird et al. 2017 for list of samples).

#### Diversity and genetic structure

To determine patterns of diversity within populations of *B. rapa*, we evaluated nucleotide diversity (π; Nei and Li, 1979) in Tassel 5 (Glaubitz et al., 2014) and variance within and across populations and geographical groups using the Analysis of Molecular Variance (AMOVA) (Excoffier et al., 1992) in Arlequin 3.5 (Excoffier et al., 2005). Given that the data are GBS derived SNPs which inherently optimize variability across individuals by targeting segregating sites, the ability to compare these nucleotide diversity results with those in other species or π calculated using other types of data are limited. For the AMOVA analysis, groups were defined geographically as follows: East Asia, Central Asia, Europe, and Americas. Six turnips of unknown provenance since they would not be informative. With the exception of the TreeMix analysis, which used populations delineated based on monophyletic clades through SNPhylo (Supp. fig. S2), populations were delineated for all analyses based on clades recovered from the RAxML tree in conjunction with phenotypic data and provenance information. The polyphyletic weedy *B. rapa* populations were delineated based on provenance (weedy Europe, weedy Americas) except for the Weedy CSI (Caucasus, Siberia, Italy) clade which consistently formed a discrete clade distinct from these other lineages.

To determine the genetic structure of our samples, we used fastSTRUCTURE 1.0 (Raj et al. 2014). The *Brassica oleracea* outgroup sample was omitted from this analysis. We ran fastSTRUCTURE at group (K) values between 1 and 20 and used the *ChooseK*.*py* script included in the fastSTRUCTURE package to assess the K value that maximized marginal likelihood. FastSTRUCTURE plots were generated using the online utility STRUCTURE PLOT 2.0 (Ramasamy et al., 2014). To further investigate the genetic structure of the samples, we used principal component analysis (PCA) in Plink 1.07 (Purcell et al., 2007), running analyses that include all populations at once as well subsets of populations focusing on crops within geographic regions. To evaluate genetic differentiation among populations, we calculated F_ST_ (Weir and Cockerham, 1984) between each pair of populations defined by subspecies and geography using Arlequin 3.5 (Excoffier et al., 2005).

#### Tree-based analyses

In order to visualize the hierarchical structure of the species, we generated a neighbor-joining tree (Saitou and Nei, 1987) using Nei’s genetic distance and 100 bootstrap replicates in PAUP 4.0 (Swofford, 2003), a maximum likelihood tree with RAxML 8.0 (Stamatakis, 2014) on the CIPRES platform (Miller at el. 2012) using rapid bootstrapping (*-fa* and *-x* flags), 100 bootstraps, a GTR+Γ substitution model, and sample NGB 162411 (*Brassica oleracea* from Denmark) as an outgroup, and SNPhylo v. 20160204 (Lee et al. 2014) was run using 0.1 for LD threshold, 0.01 for minor allele threshold, 0.1 for missing rate, 1000 bootstrap replicates, and again rooted with *B. oleracea*.

#### TreeMix and f_4_-statistics

To test species tree relationships and introgression we used TreeMix v. 1.13 (Pickrell and Pritchard 2012). To reduce signals of recent introgression, samples were pruned to include only those recovered as monophyletic clades in the SNPhylo analysis (Supp. fig. 2) and analyses were run for 0 to 8 migration edges with the following parameters: *-root B. oleracea, - k 300*. To provide additional support for inferred migration edges four-population (*f*_4_) tests (Reich et al. 2009) for treeness were used as implemented in TreeMix.

#### Demographic analysis

In order to investigate population branching structure, population size changes, and migration between populations, we constructed and tested demographic models using *moments* v1.0.1 (Jouganous et al. 2017), which uses the joint allele frequency spectrum (AFS) to fit parameterized demographic models. For this analysis, we did not filter by minimum minor allele frequency as it can artificially skew the AFS and bias inferences. Because numerous SNPs might not be called in every individual in each population, we projected sample sizes down to include more SNPs in each analysis (Marth et al. 2004). Total sample sizes and projection sizes are given in table S6. For all models, we ran parameter inference using the folded AFS (using the minor allele frequency of all biallelic SNPs) to avoid biases caused by misidentification of the ancestral allele.

We tested demographic models using between two and four populations. To reduce the risk of model over-parameterization, population sizes were allowed to vary between branches but were constant along each branch, and we allowed no more than two symmetric migration rates per model. We tested models for a total of 13 unique sets of populations (table S9). For inferences involving four populations we fixed best-fit parameters from simpler model inferences and only fit parameters for the additional populations. All models are illustrated in Supp. fig. S7, with parameter descriptions in table S7. Best-fit parameters were inferred using the built-in composite likelihood-based optimization functions in *moments*, and confidence intervals were computed using the Godambe information matrix, which corrects test statistics to account for linkage between SNPs (Coffman et al. 2016).

To clarify the relative order of population splits and the mode of divergence (e.g., with or without ongoing gene flow or via admixture events) in the evolutionary history of turnips, toria, and the newly recovered clade of weedy samples from the Caucasus, Siberia, and Italy, we tested additional demographic models. These populations were selected because the PCA, FastStructure, and tree-based analyses suggested a potential key role in the early origins of *B. rapa* crops. To try to resolve these deeper splits, we tested the following parameterized demographic models using sets of three populations including the weedy CSI lineage: 1) Weedy CSI, Central Asian turnips, and East Asian turnips, 2) Weedy CSI, toria, and Central Asian turnips, 3) Weedy CSI, toria, and European turnips, and 4) Weedy CSI, toria, and East Asian turnips. For each set of three populations, we tested models with each of the three populations as the earlier diverging branch, as well as a model in which the weedy CSI were formed through an admixture event between the other two populations which had earlier diverged. For each of these four settings, we tested models that both allow and disallow migration, so that we fit eight models for each set of three populations.

Because computing the AFS for more than four populations is computationally burdensome, we compiled the demographic model for ten populations across the Eastern and Western clades by parsimoniously combining inferred models for subsets of these populations. This combined model should be treated with caution, as parameters were not refitted over the 10-population AFS. However, the topology and approximate split times are largely consistent with the inferred models for the 13 population subsets in Table S9. *Moments* returns parameter estimates in genetic units, scaled by 2*N*_*e*_, where *N*_*e*_ is the ancestral effective population size. We used an estimate of *N*_*e*_≅40,000 from Qi et al. (2017), which assumed a mutation rate of 9e-9 (Kagale et al., 2014). To convert from generations to years, we assumed one year per generation. The parameters in all inferred demographic models using *moments* are in genetic units, i.e. scaled by the ancestral Ne, which is set to 1. Thus, all population sizes nu_pop are relative sizes in comparison to the ancestral reference Ne. Times of splits and size changes are given in units of 2Ne generations, and migration rates are given in units of 2Nemper-gen, where mper-gen is the per-generation probability that a lineage in one population had its parent in another source population. In all models, theta=4NeL is a free parameter that accounts for the linear scaling of the total number of variants in the data AFS, where is the per-generation, per-site mutation rate and L is the effective number of sequenced sites.

We computed pairwise linkage disequilibrium (LD) as 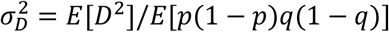 where *p* and *q* are the allele frequencies at the two loci, *D* is the standard covariance measure of LD, and expectations are estimated as averages of those statistics across pairs of loci (Ohta and Kimura, 1969).

This measure of LD is closely related to the familiar *r*^2^, but allows for unbiased comparison of LD between cohorts with varying sample size (Ragsdale & Gravel, 2020). We also computed average LD between all pairs of variants on separate chromosomes within each population, that is, average LD for unlinked loci within each population. LD between unlinked loci provides an estimate of effective population sizes over the last few generations, since expected 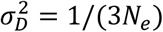 for unlinked loci in randomly mating outbred populations (Waples, 2006; Ragsdale and Gravel, 2020), and rearranging this equation provides an estimate for *N*_*e*_. Long-range LD is not only sensitive to *N*_*e*_, but can also be elevated due to recent admixture between diverged populations. However, LD between unlinked loci decays within just a handful of generations, so that *N*_*e*_ based on unlinked-LD is largely insensitive to admixture that occurred more than 2-4 generations ago (Waples, 2006).

#### Species Distribution modeling

To obtain georeferenced *B. rapa* occurrence data, we drew from the Global Biodiversity Information Facility (GBIF, http://www.gbif.org). We removed duplicate records, records without location data, records without vouchers, and records from botanical gardens, and to ensure that the occurrence data included only non-crop *B. rapa*, we included only samples with location data suggesting uncultivated status. Occurrences were included that referenced roadsides, ditches, waste areas, abandoned fields, railroad tracks, parklands, or weediness in their metadata, or were originally determined as the weedy ssp. *campestris* or ssp. *sylvestris*. Our initial GBIF search for *Brassica rapa* returned 34,361 occurrences. After filtering using the above criteria, 2263 occurrences remained. To reduce bias in sampling that may arise from collecting in accessible areas close to population centers, we removed samples occurring within 100km of each other using the spThin R package (Aiello-Lammens et al., 2015). Spatial thinning resulted in a final dataset of 638 occurrences. We obtained GIS rasters for 19 bioclimatic variables (Supp. table S10) with 2.5-minute spatial resolution from WorldClim Version 1.4 (Hijmans et al., 2005). We modeled the niche of weedy *B. rapa* and predicted compatible habitat given the CCSM4 mid-Holocene climate model (Meehl et al., 2012) and “contemporary” WorldClim data collected between 1960-1990 in MaxEnt (Phillips et al., 2006). For both contemporary and mid-Holocene models, all 19 variables were retained as recent simulations indicate that removing collinear variables does not significantly impact model performance with maximum entropy modeling (Feng et al., 2019) Model performance was evaluated with the adjusted area under receiver operating characteristic (ROC) curve (DeLong et al., 1988). The linear, quadratic, product, and hinge features were used and jackknifes were used to measure variable importance.

## Acknowledgements

This work was supported by a National Science Foundation Doctoral Dissertation Improvement Grant (grant number DEB-1601430 to A.C.M. and E.E.) and the funding from the University of Wisconsin-Madison (UW-Madison) Department of Botany. J.C.P., K.B., H.A., and M.E.M. were supported by the US Department of Energy (DOE HDTRA 1-16-1-0048), the US Department of Agriculture, and the US National Science Foundation (NSF IOS 1339156). We would like to thank the staff of UW-Madison Biotechnology Center and Bioinformatics Resource Center for their support with sequencing and analysis and the Walnut Street Greenhouse at UW-Madison for assistance with growing the plants for phenotyping.

## Footnotes

1 While the leaves and petioles of turnip crops are used for food in many cultures and leafy types without enlarged root-hypocotyls are sometimes referred to as “turnip tops” or “turnip greens” (Martínez et al. 2013), the crops referred to in this study as leafy types (e.g., rapini, grelos, bok choy, napa cabbage) do not have swollen below-ground storage organs.

## Notes

https://doi.org/10.5061/dryad.0gb5mkm0m

### Competing Interest Statement

The authors have declared no competing interest.

